# Trans-ancestral GWAS of alcohol dependence reveals common genetic underpinnings with psychiatric disorders

**DOI:** 10.1101/257311

**Authors:** Raymond K. Walters, Mark J. Adams, Amy E. Adkins, Fazil Aliev, Silviu-Alin Bacanu, Anthony Batzler, Sarah Bertelsen, Joanna Biernacka, Tim B. Bigdeli, Li-Shiun Chen, Toni-Kim Clarke, Yi-Ling Chou, Franziska Degenhardt, Anna R. Docherty, Pierre Fontanillas, Jerome Foo, Louis Fox, Josef Frank, Ina Giegling, Scott Gordon, Laura M. Hack, Annette M. Hartmann, Sarah M. Hartz, Stefanie Heilmann-Heimbach, Stefan Herms, Colin Hodgkinson, Per Hoffmann, Jouke-Jan Hottenga, Martin A. Kennedy, Mervi Alanne-Kinnunen, Bettina Konte, Jari Lahti, Marius Lahti-Pulkkinen, Lannie Ligthart, Anu-Maria Loukola, Brion S. Maher, Hamdi Mbarek, Andrew M. McIntosh, Matthew B. McQueen, Yuri Milaneschi, Teemu Palviainen, John F. Pearson, Roseann E. Peterson, Renato Polimanti, Samuli Ripatti, Euijung Ryu, Nancy L. Saccone, Jessica E. Salvatore, Sandra Sanchez-Roige, Melanie Schwandt, Richard Sherva, Fabian Streit, Jana Strohmaier, Nathaniel Thomas, Jen-Chyong Wang, Bradley T. Webb, Robbee Wedow, Leah Wetherill, Amanda G. Wills, 23andMe Research Team, Jason D. Boardman, Danfeng Chen, Doo-Sup Choi, William E. Copeland, Robert C. Culverhouse, Norbert Dahmen, Louisa Degenhardt, Benjamin W. Domingue, Sarah L. Elson, Mark Frye, Wolfgang Gäbel, Marcus Ising, Emma C. Johnson, Margaret Keyes, Falk Kiefer, John Kramer, Samuel Kuperman, Susanne Lucae, Michael T. Lynskey, Wolfgang Maier, Karl Mann, Satu Männistö, Jeanette Nance McClintick, Jacquelyn L. Meyers, Bertram Müller-Myhsok, John I. Nurnberger, Aarno Palotie, Ulrich Preuss, Katri Räikkönen, Maureen D. Reynolds, Monika Ridinger, Norbert Scherbaum, Marc Shuckit, Michael Soyka, Jens Treutlein, Stephanie Witt, Norbert Wodarz, Peter Zill, Daniel E. Adkins, Joseph M. Boden, Dorret Boomsma, Laura J Bierut, Sandra A. Brown, Kathleen K. Bucholz, Sven Cichon, E. Jane Costello, Harriet de Wit, Nancy Diazgranados, Danielle M. Dick, Johan G. Eriksson, Lindsay A. Farrer, Tatiana M. Foroud, Nathan A. Gillespie, Alison A. Goate, David Goldman, Richard A. Grucza, Dana B. Hancock, Kathleen Mullan Harris, Andrew C. Heath, Victor Hesselbrock, John K. Hewitt, Christian Hopfer, John Horwood, William Iacono, Eric O. Johnson, Jaakko A. Kaprio, Victor Karpyak, Kenneth S. Kendler, Henry R. Kranzler, Kenneth Krauter, Paul Lichtenstein, Penelope A. Lind, Matt McGue, James MacKillop, Pamela A.F. Madden, Hermine Maes, Patrik Magnusson, Nicholas G. Martin, Sarah E. Medland, Grant W. Montgomery, Elliot C. Nelson, Markus Nöthen, Abraham A. Palmer, Nancy L. Pedersen, Brenda W.J.H. Penninx, Bernice Porjesz, John P. Rice, Marcella Rietschel, Brien P. Riley, Richard Rose, Dan Rujescu, Pei-Hong Shen, Judy Silberg, Michael C. Stallings, Ralph E. Tarter, Michael M. Vanyukov, Scott Vrieze, Tamara L. Wall, John B. Whitfield, Hongyu Zhao, Benjamin M. Neale, Joel Gelernter, Howard J. Edenberg, Arpana Agrawal

## Abstract

Liability to alcohol dependence (AD) is heritable, but little is known about its complex polygenic architecture or its genetic relationship with other disorders. To discover loci associated with AD and characterize the relationship between AD and other psychiatric and behavioral outcomes, we carried out the largest GWAS to date of DSM - IV diagnosed AD. Genome - wide data on 14,904 individuals with AD and 37,944 controls from 28 case / control and family - based studies were meta - analyzed, stratified by genetic ancestry (European, N = 46,568; African; N = 6,280). Independent, genome - wide significant effects of different *ADH1B* variants were identified in European (rs1229984; p = 9.8E - 13) and African ancestries (rs2066702; p = 2.2E - 9). Significant genetic correlations were observed with schizophrenia, ADHD, depression, and use of cigarettes and cannabis. There was only modest genetic correlation with alcohol consumption and inconsistent associations with problem drinking. The genetic underpinnings of AD only partially overlap with those for alcohol consumption, underscoring the genetic distinction between pathological and non - pathological drinking behaviors.

## INTRODUCTION

Excessive alcohol use is a leading contributor to morbidity and mortality. One in 20 deaths worldwide is attributable to alcohol consumption, as is 5.1% of the global burden of disease^1^. Alcohol dependence (AD), as defined by the Fourth Edition of the American Psychiatric Association Diagnostic and Statistical Manual of Mental Disorders (DSM - IV) ^2^, is a serious psychiatric disorder characterized by tolerance, withdrawal, loss of control over drinking and excessive alcohol consumption despite negative health and social consequences. Among alcohol drinkers, 12% meet criteria for DSM - IV AD during their lifetimes^3^. In the United States, only 25% of those with AD ever receive treatment^4, 5^.

AD is moderately heritable (49% by a recent meta - analysis) ^6^ and numerous genome - wide association studies (GWAS) have aimed to identify loci contributing to this genetic variance (see^7^ for a review). According to one study, common SNPs are responsible for as much as 30% of the variance in AD^8^, but few have been identified to date. Variants in the genes responsible for alcohol metabolism^9-19^ (*ADH1B* and, to a lesser extent, *ADH1C* and others^20-22^, e.g., *ADH4)* have been strongly implicated, initially in East - Asians^9,11,12^ and more recently in people of European origin (EU) and in African - Americans (AAs) ^13-15^. The association between AD (and related problem drinking phenotypes) and rs1229984, a missense SNP (Arg48His) in *ADH1B* that affects the conversion of alcohol to acetaldehyde, represents one of the largest common - variant effect sizes observed in psychiatry, with the His48 allele accelerating ethanol metabolism and affording approximately 3 - fold reduction in likelihood of AD across numerous studies (e.g., ^14,23,24^). Another functional polymorphism, rs671 in *ALDH2* (Glu504Lys), strongly affects alcohol metabolism by blocking conversion of acetaldehyde to acetate, but is rare except in some Asian populations^9-11,17,18^. *ADH1B* and *ALDH2* polymorphisms, however, only explain a small proportion of the heritable variation in AD in populations of European ancestry.

In this study, we compiled the largest numbers of carefully diagnosed alcohol dependent individuals and alcohol - exposed controls to date, from both case - control and family studies. These included substantial numbers of both European ancestry (EU, N = 46,568, including 38,686 unrelated individuals) and African - American ancestry (AA, N = 6,280, including 5,799 unrelated individuals) subjects. Each study was subjected to stringent quality control (QC) before conducting GWAS within each population of each study, followed by a genome - wide meta - analysis. We estimated the heritability (SNP - *h*^2^) of AD and examine the extent to which aggregate genetic variation in AD is related to traits from 42 other GWAS, including continuous measures of alcohol consumption.

## METHODS

*Samples*: We collected individual genotypic data from 14 case / control studies and 9 family - based studies and summary statistics from GWAS of AD from 5 additional cohorts (**Table 1**; see **Supplementary Information** for cohort descriptions). AD was defined as meeting criteria for a DSM - IV (or DSM - IIIR in one instance) diagnosis of AD. Excepting three cohorts with population - based controls (N = 7,015), all controls were screened for AD. Individuals with no history of drinking alcohol and those meeting criteria for DSM - IV alcohol abuse were additionally excluded as controls where applicable (**Supplementary Information**).

**Table 1:** Descriptive statistics for cohorts in the meta - analysis of AD.

| Dataset | PMID | Male (%) | Ages (years) | European (EU) |  |  |  | African - American (AA) |  |  |  |
| --- | --- | --- | --- | --- | --- | --- | --- | --- | --- | --- | --- |
|  |  |  |  | N Total |  | N Unrelated |  | N Total |  | N Unrelated |  |
|  |  |  |  | Case | Control | Case | Control | Case | Control | Case | Control |
| Case-control: Logistic Regression |  |  |  |  |  |  |  |  |  |  |  |
| Comorbidity and Trauma Study (CATS) | 23303482 | 56% | 18-67 | 572 | 817 | 572 | 817 | -- | -- | -- | -- |
| Christchurch Health and Development Study (CHDS) | 23255320 | 48% | 16-30 | 112 | 500 | 112 | 500 | -- | -- | -- | -- |
| Collaborative Study of the Genetics of Alcoholism - case-control cohort (COGA-cc) | 20201924 | 54% | 18-79 | 583 | 363 | 583 | 363 | -- | -- | -- | -- |
| Family Study of Cocaine Dependence (FSCD) | 18243582 | 51% | 18-60 | 266 | 174 | 266 | 174 | 255 | 241 | 255 | 241 |
| German Study of the Genetics of Alcoholism (GESGA) | 19581569 | 65% | 18-84 | 1314 | 2142 | 1314 | 2142 | -- | -- | -- | -- |
| Gene-Environment Development Initiative - Great Smoky Mountains Study (GEDI-GSMS) | 8956679 | 57% | 9-26 | 42 | 565 | 42 | 565 | -- | -- | -- | -- |
| Center on Antisocial Drug Dependence (CADD) | 25637581 | 70% | 13-20 | 400 | 577 | 400 | 577 | 51 | 51 | 51 | 51 |
| Phenomics and Genomics Sample (PAGES) | 28371232 | 57% | 18-74 | 37 | 523 | 37 | 523 | -- | -- | -- | -- |
| Collaborative Study on the Genetics of Nicotine Dependence (COGEND Nico) | 17158188 | 34% | 25-82 | 135 | 272 | 135 | 272 | 46 | 232 | 46 | 232 |
| COGEND - Study of Addiction: Genetics and Environment (COGEND SAGE) | 20202923 | 37% | 18-77 | 311 | 225 | 311 | 225 | 104 | 103 | 104 | 103 |
| Spit For Science | 24639683 | 36% | >18 | 252 | 1863 | 252 | 1863 | 74 | 841 | 74 | 841 |
| National Institute on Alcohol Abuse and Alcoholism Intramural (NIAAA) | n/a | 67% | >18 | 442 | 206 | 442 | 206 | 404 | 110 | 404 | 110 |
| Mayo Clinic Center for the Individual Treatment of Alcohol Dependence (CITA) | 25290263 | 55% | ≥18 | 378 | 646 | 378 | 646 | -- | -- | -- | -- |
| Alcohol Dependence in African Americans (ADAA) | n/a | 57% | 18-69 | -- | -- | -- | -- | 794 | 297 | 794 | 297 |
| Family-based, twins and sibs: GEE |  |  |  |  |  |  |  |  |  |  |  |
| Brisbane Longitudinal Twin Study (BLTS) | 23187020 | 43% | 18-30 | 60 | 938 | 51 | 546 | -- | -- | -- | -- |
| GEDI - Virginia Twin Study on Adolescent Behavioral Development (GEDI-VTSABD) | 9294370 | 38% | 8-32 | 209 | 503 | 188 | 318 | -- | -- | -- | -- |
| Minnesota Center for Twin and Family Research (MCTFR) | 23942779 | 41% | 16-21 | 609 | 2100 | 553 | 1274 | -- | -- | -- | -- |
| Center for Education and Drug Abuse Research (CEDAR) | 21514569 | 63% | 16-34 | 59 | 200 | 54 | 152 | -- | -- | -- | -- |
| Swedish Twin Registry (STR) | 23137839 | 47% | 40-83 | 76 | 8311 | 76 | 6112 | -- | -- | -- | -- |
| Yale-Penn | 24166409 | 58% | 16-79 | 1094 | 301 | 1004 | 252 | -- | -- | -- | -- |
| Family-based, large/complex pedigrees: Logistic Mixed Model |  |  |  |  |  |  |  |  |  |  |  |
| Collaborative Study of the Genetics of Alcoholism - family cohort (COGA-fam) | 23089632 | 45% | 12-88 | 605 | 682 | 168 | 138 | -- | -- | -- | -- |
| Australian Alcohol and Nicotine Studies (OZ-ALC-NAG) | 21529783 | 45% | 18-82 | 1571 | 3069 | 1111 | 805 | -- | -- | -- | -- |
| Irish Affected Sib Pair Study of Alcohol Dependence (IASPSAD) | 15770118 | 50% | 17-84 | 721 | 1814 | 436 | 1802 | -- | -- | -- | -- |
| Yale-Penn | 24166409 | 51% | 16-79 | -- | -- | -- | -- | 1607 | 1070 | 1263 | 933 |
| Summary statistics |  |  |  |  |  |  |  |  |  |  |  |
| Netherlands Study of Depression and Anxiety / Netherlands Twin Registry (NESDA/NTR) | 18197199 | 31% | >18 | 390 | 1633 | 390 | 1633 | -- | -- | -- | -- |
| Finnish Nicotine Addiction Genetics Project (NAG-Fin) | 17436240 | 52% | 30-92 | 439 | 1137 | 439 | 1137 | -- | -- | -- | -- |
| FinnTwin12 (FT12) | 17254406 | 47% | 20-27 | 88 | 874 | 88 | 874 | -- | -- | -- | -- |
| National Longitudinal Study of Adolescent to Adult Health (Add Health) | 25378290 | 47% | 24-34 | 768 | 2981 | 768 | 2981 | -- | -- | -- | -- |
| Helsinki Birth Cohort Study (HBCS) | 16251536 | 43% | 56-70 | 36 | 1583 | 36 | 1583 | -- | -- | -- | -- |
| Total |  |  |  | 11569 | 34999 | 10206 | 28480 | 3335 | 2945 | 2991 | 2808 |

### Quality control and imputation

Data for the genotyped cohorts that shared raw data were deposited to a secure server for uniform quality control (QC). QC and imputation of the 14 case / control studies was performed using the ricopili pipeline (https://github.com/Nealelab/ricopili). For 9 family - based cohorts, an equivalent pipeline, picopili (https://github.com/Nealelab/picopili), was developed for QC, imputation, and analysis appropriate for diverse family structures, including twins, sibships and extended pedigrees (**Supplementary Information**).

After initial sample and variant QC, principal components analysis (PCA) was used to identify population outliers for exclusion and to stratify samples in each study by continental ancestry. Identified EU and AA ancestry populations were confirmed by PCA with the 1000 Genomes reference panel^25^. Final sample and variant QC, including filters for call rate, heterozygosity, and departure from Hardy - Weinberg equilibrium (HWE), was then performed within each ancestry group in each cohort (see **Supplementary Information**). Samples were also filtered for cryptic relatedness within and between cohorts and for departures from reported pedigree structures.

Each cohort was imputed using SHAPEIT^26^ and IMPUTE2^27^, using the cosmopolitan (all ancestries) 1000 Genomes reference panel. Consistency of minor allele frequencies (MAF) with the reference panel was verified prior to imputation, with SNPs in EU cohorts compared to MAF in European population samples and AA cohorts compared to MAF in African population samples. Imputed SNPs were then filtered for INFO score > 0.8 and allele frequency > 0.01 prior to analysis.

### Association Analysis

A GWAS for AD status was performed within each ancestry stratum of each sample using an association model appropriate for the study design (**Table 1**). For case / control studies, GWAS was performed using logistic regression with imputed dosages. For family - based studies of small, simple pedigrees (e.g., sibships), association with imputed genotypes was tested using generalized estimating equations (GEE). For more complex pedigrees, imputed genotypes were tested using logistic mixed models. Sex was included as a covariate, along with principal components to control for population structure (**Supplementary Information**). Details of the analytic model, software used, effective N, number of SNPs and principal components are presented for each sample in Supplementary **Table S1**.

In addition to this primary analysis, subsets of genetically unrelated individuals were selected from each family - based cohort (i.e. taking one individual per family) and used to perform a conventional case / control GWAS using logistic regression. This was used in place of the family - based GWAS for estimation of effect sizes and inclusion in estimation of SNP - *h*^2^ and genetic correlations (*r*_*g*_) using LD score regression analyses.

### Meta - analysis

The primary discovery meta - analysis of all ancestry - stratified GWAS (N_case_ = 14,904; N_control_ = 37,994) was conducted in METAL^28^. As the different study designs (family vs. case - control) produced effect sizes that were not comparable, results were combined using weighting by effective sample size (see **Supplementary Information**). Separate ancestry - specific discovery meta - analyses of EU (N = 46,568) and AA (N = 6,280) cohorts, respectively, were also performed. Heterogeneity was evaluated across all cohorts and between study design subsets (**Supplementary Information**). Power analysis was performed using CaTS^29^ with the estimated effective sample size.

In addition to the discovery meta - analyses, we conducted meta - analyses for two design subsets. First, we performed sample size weighted meta - analysis of the subset of genetically unrelated individuals in EU (N = 38,686) and AA (N = 5,799) cohorts for use in LD score regression (LDSR) analysis. Second, we performed inverse - variance weighted meta - analysis of genetically unrelated individuals in genotyped cohorts to estimate within - ancestry effect sizes for EU (N = 28,757) and AA (N = 5,799). These effect sizes were then used to compare trans - ancestral fine mapping results using inverse - variance weighted fixed effects, random effects^30^, and Bayesian^31^ models (**Supplementary Information**). Supplementary **Table S2** provides an overview of the various meta - analytic models that were fitted to data.

### Heritability and Genetic Correlation Analysis

LDSR analysis^32^ was performed to estimate the heritability explained by common SNPs in meta - analyses of unrelated EU and AA samples, respectively. LDSR was performed using HapMap3 SNPs and LD scores computed from 1000 Genomes reference samples corresponding to each population (Supplementary Information). Conversion of *h*^2^ estimates from observed to liability scale was performed assuming population prevalences of 0.159 and 0.111 for AD in alcohol - exposed EU and AA individuals, respectively^3^.

Genetic correlation between AD and 42 traits from LD Hub^33^ and other published studies^34-44^ was examined with the same unrelated EU meta - analysis (10,206 cases and 28,480 controls) and precomputed European LD scores using LDSR. To avoid increasing the multiple testing burden, redundant or highly - correlated phenotypes were reduced by manually selecting the version of the phenotype with the greatest predicted relevance to AD, largest sample size, or highest heritability (**Supplementary Information**).

### Replication

As described below, a locus on chromosome 3 was genome - wide significant (GWS) in the trans - ancestral discovery meta - analysis. The minor allele, associated with lower AD risk in our analysis, had low frequency in all EU samples except the Finnish cohorts; it was also higher in AAs. To seek replication, we examined the association between this locus and DSM - IV AD in two independent AA samples (Yale - Penn 2, n = 911 cases and 599 controls, and COGA AAfGWAS, n = 880 cases and 1,814 controls; Supplementary Information) using GEE (Yale - Penn) and Genome - Wide Association / Interaction Analysis and Rare Variant Analysis with Family Data (GWAF; in COGA) respectively. Association with AD status, broadly defined using hospital and death records, was also examined in the FINRISK cohort (1,232 cases and 22,614 controls) using Firth logistic regression^45^.

## RESULTS

**GWAS meta - analyses**: In both the EU and AA analyses, GWS loci (p < 5E - 8) were identified in the ADH gene cluster on chromosome 4 (**Figure 1** for Manhattan plot; **Table 2** for top loci; Supplementary **Figure S1** for QQ plot for discovery GWAS showing polygenic signal). Examining individual populations, rs1229984 in *ADH1B* was the strongest associated signal from the analysis in EU (p = 9.8E - 13), while rs2066702, also in *ADH1B*, was the most significant variant in AA (p = 2.2E - 9; **Figure 2** shows the regional association plots for the *ADH1B* locus for the discovery, EU, AA and trans - ancestral meta - analysis). Clumping for linkage disequilibrium (LD; *r*^2^ < .1 within 500kb) suggested multiple independent signals within this locus in both populations (**Table 2**), with differing leading alleles reflecting different LD structures and allele frequencies in each population (Supplementary **Figure S2A** and **S2B** show LD patterns in the ADH locus, including *ADH1B*, in AA and EU respectively). Conditional analysis controlling for rs2066702 (Supplementary **Figure S3** for results in AA) and rs1229984 (Supplementary **Figure S4** for results in EU) was inconclusive due to limited power, but was tentatively consistent with the existence of additional independent effects in the region (Supplementary **Table S3** shows marginal and conditional effect sizes for genome - wide significant SNPs in the *ADH1B* locus). The most promising support for an independent signal arises from rs894368 (marginal odds ratio = 0.887, p = 6.9E - 7; conditional odds ratio = 0.890, p = 6.8E - 6; Supplementary Information). Results from the trans - ancestral meta - analysis reinforced the robust effects of rs1229984 and other *ADH1B* SNPs on liability to AD (regional association plot for rs1229984 in Supplementary **Figure S5A** (inverse - variance weighted), **S5B** (modified random - effects) and **S5C** (Bayesian)) across various analytic models.

**Table 2:** Top 10 loci from the discovery meta - analysis of alcohol dependence by ancestry

| SNP | CHR | BP | A1 | A2 | Gene | A1 Allele Freq. |  | INFO score |  | Effect size |  | Discovery meta-analysis p-value |  |  |
| --- | --- | --- | --- | --- | --- | --- | --- | --- | --- | --- | --- | --- | --- | --- |
|  |  |  |  |  |  | EU | AA | EU | AA | EU OR | AA OR | EU | AA | Trans |
| Trans-ancestral meta-analysis (14,904 cases, 37,944 controls) |  |  |  |  |  |  |  |  |  |  |  |  |  |  |
| rs1229984 | 4 | 100239319 | T | C | ADH1B | 0.040 | 0.014 | 0.904 | 0.910 | 0.486 | 0.912 | 9.79E-13 | 3.48E-01 | 2.18E-11 |
| rs1789912 | 4 | 100263942 | T | C | ADH1C | 0.418 | 0.132 | 1.000 | 1.020 | 1.106 | 1.211 | 1.98E-07 | 1.32E-03 | 1.47E-09 |
| rs2066702 | 4 | 100229017 | A | G | ADH1B | -- | 0.215 | -- | 0.989 | -- | 0.731 | -- | 2.21E-09 | 2.21E-09 |
| rs6827898 | 4 | 100295863 | A | G | -- | 0.123 | 0.112 | 0.963 | 0.942 | 1.145 | 1.270 | 5.21E-07 | 9.31E-04 | 2.97E-09 |
| rs7644567 | 3 | 29201672 | A | G | RBMS3 | -- | 0.705 | -- | 0.997 | -- | 1.229 | -- | 6.64E-06 | 1.36E-08 |
| rs894368 | 4 | 100309313 | A | C | -- | 0.309 | 0.386 | 0.994 | 0.962 | 0.887 | 0.981 | 1.93E-08 | 9.73E-01 | 3.30E-07 |
| rs116338421 | 8 | 145761256 | C | G | ARHGAP39 | -- | 0.172 | -- | 0.974 | -- | 0.755 | -- | 4.86E-07 | 4.86E-07 |
| rs79171978 | 12 | 17798824 | C | G | -- | 0.099 | 0.027 | 0.989 | 0.986 | 1.201 | 1.016 | 5.47E-08 | 8.18E-01 | 5.98E-07 |
| rs2461618 | 7 | 68667233 | A | G | -- | -- | 0.088 | -- | 0.984 | -- | 0.669 | -- | 6.30E-07 | 6.30E-07 |
| rs8017647 | 14 | 32456358 | T | C | -- | 0.792 | 0.565 | 0.998 | 0.991 | 0.901 | 0.923 | 8.05E-06 | 4.71E-02 | 1.03E-06 |
| African ancestry meta-analysis (3,335 cases, 2,945 controls) |  |  |  |  |  |  |  |  |  |  |  |  |  |  |
| rs2066702 | 4 | 100229017 | A | G | ADH1B | -- | 0.215 | -- | 0.989 | -- | 0.731 | -- | 2.21E-09 | 2.21E-09 |
| rs5781337 | 1 | 223883425 | CA | C | -- | 0.263 | 0.212 | 0.982 | 0.927 | 1.007 | 0.664 | 8.85E-01 | 1.62E-07 | 6.59E-02 |
| rs116338421 | 8 | 145761256 | C | G | ARHGAP39 | -- | 0.172 | -- | 0.974 | -- | 0.755 | -- | 4.86E-07 | 4.86E-07 |
| rs3857224 | 4 | 100129685 | T | C | ADH6 | 0.315 | 0.585 | 0.994 | 1.000 | 0.970 | 0.814 | 2.40E-01 | 5.86E-07 | 2.36E-03 |
| rs2461618 | 7 | 68667233 | A | G | -- | -- | 0.088 | -- | 0.984 | -- | 0.669 | -- | 6.30E-07 | 6.30E-07 |
| rs10784244 | 12 | 62035165 | G | A | -- | 0.153 | 0.484 | 1.000 | 0.998 | 1.041 | 1.226 | 6.26E-02 | 1.04E-06 | 2.49E-04 |
| rs17199739 | 16 | 25444288 | G | A | -- | 0.176 | 0.096 | 0.993 | 0.955 | 0.994 | 0.693 | 4.25E-01 | 1.11E-06 | 8.66E-03 |
| rs79016499 | 11 | 93010988 | T | C | -- | -- | 0.066 | -- | 0.928 | -- | 1.729 | -- | 1.36E-06 | -- |
| rs740793 | 17 | 3846353 | G | A | ATP2A3 | 0.453 | 0.350 | 0.973 | 0.970 | 0.996 | 1.370 | 4.66E-01 | 1.48E-06 | 3.44E-01 |
| rs143258048 | 3 | 75982870 | A | AC | ROBO2 | -- | 0.028 | -- | 0.879 | -- | 0.490 | -- | 1.86E-06 | -- |
| European ancestry meta-analysis (11,569 cases, 34,999 controls) |  |  |  |  |  |  |  |  |  |  |  |  |  |  |
| rs1229984 | 4 | 100239319 | T | C | ADH1B | 0.040 | 0.014 | 0.904 | 0.910 | 0.486 | 0.912 | 9.79E-13 | 3.48E-01 | 2.18E-11 |
| rs894368 | 4 | 100309313 | A | C | -- | 0.309 | 0.386 | 0.994 | 0.962 | 0.887 | 0.981 | 1.93E-08 | 9.73E-01 | 3.30E-07 |
| rs3811802 | 4 | 100244221 | G | A | ADH1B | 0.454 | 0.529 | 0.958 | 0.956 | 1.162 | 0.914 | 2.40E-08 | 2.19E-02 | 1.22E-04 |
| rs79171978 | 12 | 17798824 | C | G | -- | 0.099 | 0.027 | 0.989 | 0.986 | 1.201 | 1.016 | 5.47E-08 | 8.18E-01 | 5.98E-07 |
| rs1154445 | 4 | 100288521 | G | T | -- | 0.425 | 0.134 | 0.970 | 0.986 | 1.137 | 1.211 | 1.80E-07 | 2.63E-02 | 1.48E-08 |
| rs6827898 | 4 | 100295863 | A | G | -- | 0.123 | 0.112 | 0.963 | 0.942 | 1.145 | 1.270 | 5.21E-07 | 9.31E-04 | 2.97E-09 |
| rs4388946 | 12 | 17935154 | C | A | -- | 0.240 | 0.297 | 0.988 | 0.976 | 1.137 | 0.950 | 7.14E-07 | 1.87E-01 | 7.05E-05 |
| rs1229863 | 4 | 100252386 | A | T | ADH1B | 0.174 | 0.038 | 0.990 | 0.989 | 1.145 | 1.254 | 7.80E-07 | 4.26E-02 | 9.28E-08 |
| rs34929220 | 15 | 69769635 | T | C | DRAIC | 0.690 | 0.937 | 0.898 | 0.943 | 0.893 | 1.028 | 1.02E-06 | 8.38E-01 | 7.38E-06 |
| rs113659074 | 4 | 100252308 | T | G | ADH1B | 0.068 | 0.093 | 0.980 | 0.947 | 0.800 | 1.166 | 1.54E-06 | 6.63E-02 | 2.99E-04 |

**Figure 1:**
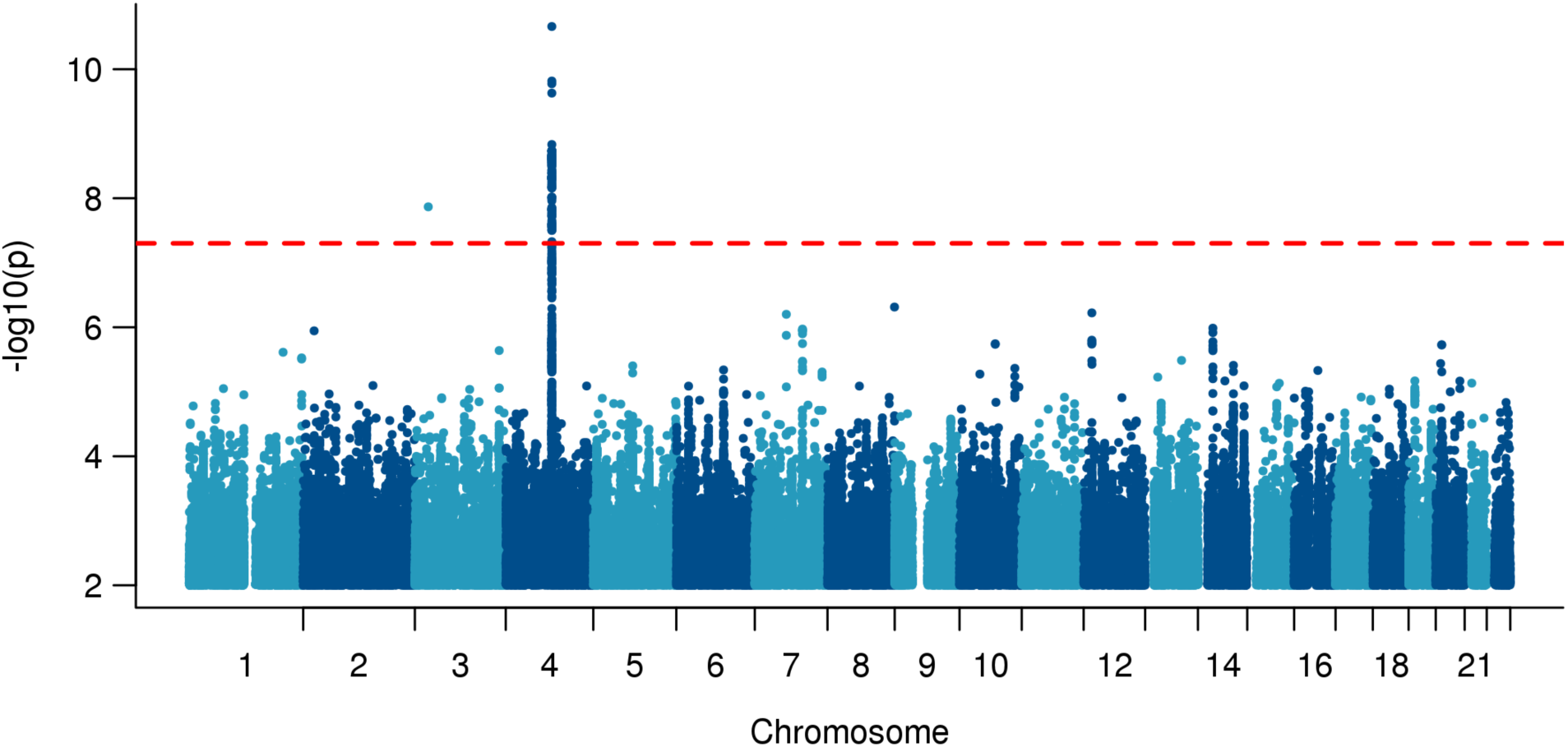
Manhattan plot of discovery trans - ancestral meta - analysis showing strong evidence for rs1229984 in *ADH1B*.

**Figure 2:**
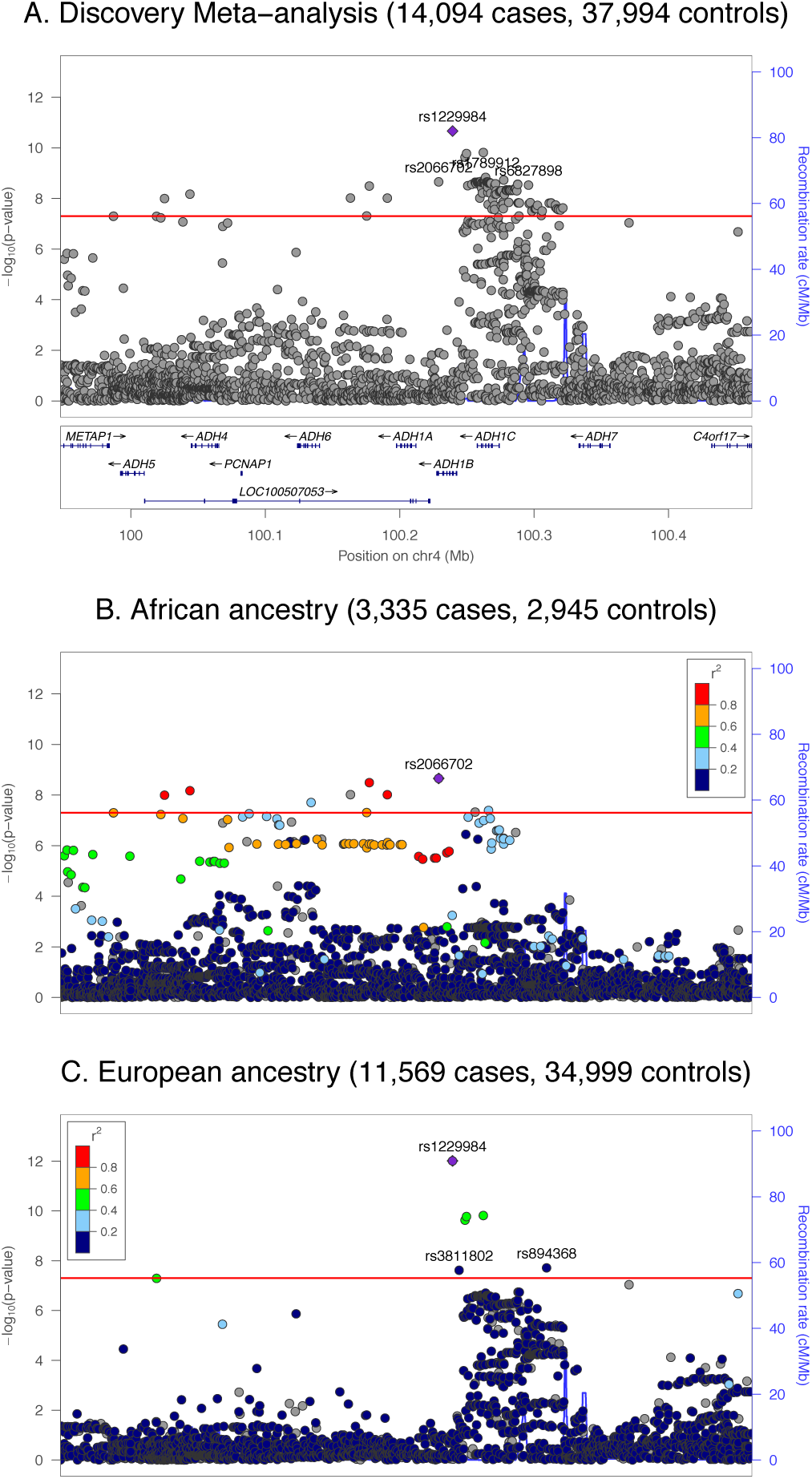
Regional plots for the ADH1B locus, rs1229984, in the European (EU), African - American (AA) and trans - ancestral discovery meta - analysis.

We also verified whether variants affecting *ADH1B* expression (eQTLs) were associated with AD. Considering GTEx data V7 (available at http://www.gtexportal.org/), 263 variants were reported to affect *ADH1B* expression in different human tissues (FDR q<0.05). After LD - informed clumping and the exclusion of variants in LD with the GWS coding alleles (i.e., rs1229984 and rs2066702), three variants (i.e., rs11939328, rs10516440, rs7664780) were considered with respect to their association with AD. SNP rs10516440 showed a genome - wide significant association with AD with contribution from both AA and EU analyses (trans - ancestry p = 4.72E - 8; EU p = 3.97E - 6; AA p = 1.97E - 3). In line with the effect of the coding variants where the protective allele is associated with increased *ADH1B* enzymatic activity, the rs10516440*A allele was associated with reduced AD risk and increased *ADH1B* expression, which was consistent across multiple tissues (multi - tissue p = 1.42E - 76).

A novel locus on chromosome 3, rs7644567, also achieved GWS in the meta - analysis (p = 3.03E - 8; Supplementary **Figure S6** for regional association plot), primarily attributable to contributions from the AA samples (p = 6.64E - 6) with the major, A, allele being associated with AD risk liability. The G allele has an MAF = 0.29 in AA, but MAF<0.01 in most EU samples, except in FinnTwin (MAF = .032) and NAG - Fin (MAF = 0.054). In AA, rs7644567 does not appear to be in high LD with other variants (Supplementary **Figure S6**) and did not replicate in two independent AA samples. In the independent FINRISK cohort (MAF = .045), there was modest evidence for association (p = 0.019), but with risk associated with the minor G allele (**Supplementary Table S4** for results in each replication sample).

Overall, there was limited evidence for heterogeneity across all cohorts, within ancestry, between ancestries, or between study designs within ancestry (**Supplementary Information**; Supplementary **Figure S7 - S13**). Gene - level association testing with MAGMA^46^ did not identify any additional genes in EU or AA (Supplementary **Table S5** for top 20 genes in EU and AA).

**Heritability and genetic correlations:** LD score based liability - scale SNP - heritability of AD was estimated at *h*^2^_*g*_ = 0.090 (SE = 0.019, p = 8.02E - 7) in the meta - analysis of unrelated EU samples. Exclusion of the *ADH1B* locus did not substantially modify this estimate (*h*^2^_*g*_ = 0.089, SE = 0.0185). Nominally significant heritability from common variants was also estimated for the meta - analysis of unrelated AA individuals based on LDSR with scores computed from 1000 Genomes African populations (p = .017), but the quantitative estimate of *h*^2^_*g*_ was unstable depending on the choice of reference panel, reflecting the challenge of correctly specifying LDSR and robustly modelling LD for the admixed AA population (**Supplementary Information**).

Significant genetic correlation with AD in EU was observed for 16 traits (significant genetic correlations in **Figure 3;** all genetic correlations in Supplementary **Table S6**), after correction for multiple testing (p = 1.19E - 3 for 42 traits). The largest positive correlations were with ever smoking tobacco (rg = .708, p = 1.3E - 7) and lifetime cannabis use (rg = .793, p = 2.5E - 4), and with other psychiatric disorders and traits, especially schizophrenia (rg = 0.357, p = 3.2E - 11), ADHD (rg = .444, p = 4.2E - 6), and depressive symptoms (rg = .603, p = 2.6E - 7). Educational attainment (rg = - 0.424, p = 6.8E - 9) and age at first birth (higher values indicate that subjects were older when they had their first child, rg = - 0.63, p = 2.0E - 9) showed significant inverse genetic correlation with AD suggesting that liability to AD risk was genetically related to lower educational attainment and lower age at which one had their first child.

Unexpected patterns of genetic correlation were observed when comparisons were made to other alcohol - related measures. AD was genetically correlated with alcohol consumption in a meta - analysis of the Alcohol Genome - wide Association (AlcGen) and Cohorts for Aging and Research in Genomic Epidemiology Plus (CHARGE +) consortia^36^ (rg = .695, p = 6.9E - 6) but only modestly with alcohol consumption from the recent large UK Biobank analysis^37^ (rg = 0.371, p = 5.2E - 5). Liability to AD was not correlated with genome - wide SNPs from a recent GWAS of the Alcohol Use Disorders Identification Test (AUDIT) in 23andMe^38^ (rg = 0.076, p = 0.65), perhaps due to the low levels of drinking observed in this population^38^. Additional analysis indicates AD is genetically correlated with GWAS of delay discounting in the 23andMe sample^42^ (rg = 0.478, p = 6.0E - 3), suggesting behavioral phenotypes in the cohort are still informative to AD.

**Associations with other GWS loci**: We examined results for the eight independent variants associated at GWS levels with alcohol consumption in the UK Biobank^37^ (**Supplementary Table S7**). Among the UK Biobank findings, three of the four reported variants in the ADH region of chromosome 4 (rs145452708 – a proxy for rs1229984, rs29001570 and rs35081954) were associated in the present study with AD (p ranging from 3.5E - 5 – 2.3E - 10) with sign concordant effects; the remaining variant was excluded from our analysis due to MAF <0.01. The UK Biobank lead variant in *KLB*, rs11940694, was nominally associated with AD (p = .0097), though this does not surpass multiple testing correction for the eight GWS alcohol consumption loci. We see little evidence (p >0.2) for association of AD with the reported loci at *GCKR* and *CADM2*, which may be due to differences in power for the given effect size or because these genes exert an influence on liability to consume alcohol but not later problems.

The locus on chromosome 18 showed limited regional association with AD, but the index variant was not present in our analysis because it no longer appears in the 1000 Genomes Phase 3 reference panel^25^.

**Power analysis:** Only 3 additional loci reach p < 1E - 6 (**Table 2**). Power analyses indicated that the current meta - analysis is expected to have at least 63% power to detect very common variants (MAF ≥ 0.25) with odds ratios ≥ 1.10 at p < 1E - 6 (41% for p < 5E - 8; **Supplementary Figure S14** for power analysis curves). Power is lower for less common variants (MAF ≥ .05) even with odds ratios ≥ 1.20 at p < 1E - 6 (60% power) and p < 5E - 8 (38% power).

## DISCUSSION

To our knowledge, this is the largest GWAS of rigorously - defined AD. We identified loci in *ADH1B* that differed between EU and AA, as well as novel genetic correlations between AD and psychiatric disorders (e.g., schizophrenia), tobacco and cannabis use, and behavioral outcomes (e.g., educational attainment). Analyses also revealed a genetic distinction between GWAS results for alcohol consumption and AD. Although larger sample sizes can be amassed by focusing on quantitative measures of consumption, only the upper tail is relevant to AD (as a medical diagnosis) and even that does not capture other aspects of disordered drinking (e.g., loss of control, withdrawal) directly. Conversely, cases derived from electronic medical records (e.g., ICD codes) may result in a high rate of false negatives, while self - screening instruments (e.g. AUDIT scores) is best suited to analyses of disordered drinking when a sufficiently high threshold or score cut - off is applied to pinpoint severity. Our study has the advantage of greater diagnostic precision via use of semi - structured interviews to diagnose AD systematically in a majority of the constituent studies.

The genome - wide significant SNPs reaffirm the importance of functional variants affecting alcohol metabolism to the risk of AD. The top association in *ADH1B*, rs1229984, is a missense variant that is amongst the most widely studied in relation to alcohol use, misuse and dependence. The resulting amino acid substitution (Arg48His) increases the rate at which ADH1B oxidizes ethanol to acetaldehyde^10, 11^. Early studies on Asian populations in which the derived allele is common demonstrated strong protection against the development of AD^9-11^. In EUs and AAs, the protective allele is present at much lower frequencies (EU MAF = 3 - 4%, AA MAF < 1%), but recent large - scale studies have shown an association between this locus and alcohol consumption and problems at GWS levels in EU with similar effect size^14, 15^. The lead variant in AA cohorts, rs2066702 (Arg370Cys), is another functional missense variant in *ADH1B*, and it, similarly, encodes an enzyme with an increased rate of ethanol oxidation^10, 11^. The allele encoding Cys370 is common among AAs, but rare in other populations^10^. Our results clearly show that these two different functional SNPs in *ADH1B* both affect risk for alcoholism, with their relative importance dependent upon allele frequency in the population studied. Larger future studies will be needed to evaluate the evidence for additional independent effects in the chromosome 4 locus.

The only other locus to reach significance was rs7644567 on chromosome 3, primarily driven by AA cohorts due to the variant’s very low MAF in EU. This locus did not replicate in independent African or Finnish ancestry samples. We note that the conventional genome - wide significance threshold is derived for European ancestry samples, and thus is likely to be too lenient in GWAS of African - ancestry cohorts due to higher genetic diversity and corresponding increase in the effective number of independent tests in the GWAS^47, 48^. As an illustration, in 4 samples from the current study that included both EU and AA participants, the number of independent SNPs identified upon LD pruning was 1.7 - to 2.3 - fold greater in AA than EU subjects. Much larger studies in AA and other non - EU populations will clearly be important to elucidate additional loci.

Despite limited SNP - level findings, there is significant evidence for polygenic effects of common variants in both EU and AA cohorts. The estimated h^2^_g_ = .09 for AD in EU is only modestly lower than those recently reported for alcohol consumption (h^2^_g_ = .13)^37^ and AUDIT scores (h^2^_g_ = .12)^38^ and comparable to estimates derived for cigarettes - per - day^33^. Our h^2^_g_ estimate is lower than a prior report^8^, likely reflecting a combination of differences in estimation method and greater heterogeneity in ascertainment strategy across samples in the current study. The latter is especially relevant given that we incorporated population - based cohorts with a wide range of ages at ascertainment and cultural environments, as well as cohorts enriched for other substance use disorders.

Comparing our GWAS to recent GWAS of alcohol consumption measures suggests that the liability underlying normative patterns of alcohol intake and AD are only partially overlapping. Genome - wide, we observe only modest genetic correlation (significantly < with log - scaled alcohol consumption by participants in AlcGen and CHARGE + Consortia cohorts^36^ (rg = .695) and in the UK Biobank^37^ (rg = .371), and no significant correlation with GWAS of log - scaled AUDIT scores in 23andMe participants^38^ (rg = .076). We also observe only partial replication of the 8 loci significantly associated with consumption in the UK Biobank. One key factor in interpreting the differences between these traits and AD is that the distribution of consumption levels and AUDIT scores can be highly skewed in population samples, with most individuals at the low (nonpathological) end of the spectrum. This effect may be especially pronounced among the older, healthy volunteers of the UK Biobank cohort^49^ and the 23andMe cohort, which is more educated and has higher socioeconomic status than the general US population ^38^. We hypothesize that the variants that affect consumption at lower levels may differ substantively from those that affect very high levels of consumption in alcohol dependent individuals, who are also characterized by loss of control over intake^50^. This appears to be the case in one prior study that used specific cut - offs to harmonize AUDIT scores with AD data and noted significant concordance in SNP - *h*^2^ estimates^51^ – according to that study, the optimal cutoffs for their sample were ≥6 and ≥9 for women and men respectively. However, there is a need for a further detailed characterization of how AUDIT cut - offs may be applied to maximize concordance with genetic liability to AD diagnosis risk. The strongly negative genetic correlation between educational attainment and AD, in contrast to positive genetic correlations of education with consumption and AUDIT scores, further underscore this distinction between normative / habitual levels of alcohol intake and diagnosed AD, at least in the respective populations studied.

The current analysis also identified robust genetic correlation of AD with a broad variety of psychiatric outcomes. This correlation is strongest for aspects of negative mood, including neuroticism and major depressive disorder, as also seen in twin studies^52, 53^ and through recent specific molecular evidence for pleiotropy^54, 55^. Taken together with evidence from other recent genomic studies^54^, and null correlations for other GWAS of alcohol consumption, these findings suggest that major depression may only share genetic liability with alcohol use at pathological levels.

AD was also negatively genetically correlated with AFB, which is an indicator of reproductive tempo and correlated with age at first consensual sexual intercourse^56^. This is consistent with evidence of common genetic liability to early, risky behaviors underlying AD and AFB^57^. Nominally significant genetic correlation with delay discounting (i.e. favoring immediate rewards) and the strong genetic correlation of AD with ADHD, cigarette smoking and cannabis use may similarly reflect a shared genetic factor for risk - taking and reduced impulse control.

Lower genetic correlations were observed for most biomedical and anthropometric outcomes. Liver enzymes GGT and ALT, once proposed as possible biomarkers for alcohol abuse^58^, showed, as expected, nominal evidence for genetic correlation with AD but neither survived multiple testing correction. Notably, we did not find any association between AD and body - mass index (BMI). Negative genetic correlations with BMI were previously reported for both alcohol consumption^37^ and AUDIT scores^38^, but there is prior evidence that BMI has differing underlying genetic architecture in the context of AD and outside of that context^59^. The negative genetic correlations observed in those studies are consistent with studies of light to moderate drinking, which is also associated with healthier lifestyle behaviors, while heavy and problematic drinking is typically associated with weight gain^60^.

This study benefits from precision in diagnostic assessment of AD, known alcohol exposure in a majority of the controls, and careful quality control that excluded overlaps of individuals between studies while combining case - control and twin / family - based study designs. Despite these strengths our sample size was insufficient to identify additional GWS loci robustly. Power analyses indicate that additional SNPs associated with AD are likely to have small effect sizes, consistent with other psychiatric disorders (e.g. depression^61^). This supports the pressing need for collection of large numbers of well characterized cases and controls. The differences between our results and the study of AUDIT scores^38^, however, highlight that ascertainment and trait definition must also be taken into account. Careful study of how screening tools, such as the AUDIT, correlate to genetic liability to AD (as defined by DSM - IV or similar) could substantially boost sample sizes for future AD GWAS. There is also a continued need to characterize the genetic architecture of AD in non - EU populations.

We show a novel genetic distinction between drinking in the pathological range (AD) and habitual drinking that does not cross the threshold into pathology or dependence. Larger future samples will allow us to uncover additional pleiotropy between pathological and non - pathological alcohol use as well as between AD and other neuropsychiatric disorders.

Overview of numbers of alcohol dependent cases and controls from each cohort in the current analysis, including the number of genetically unrelated individuals. Cohorts are listed by study design. Sample sizes are listed after QC exclusions and stratified by ancestry group. PubMed identifiers (PMID) are listed for previous publications describing each cohort, along with the percentage of male samples and the age range in the cohort.

Top 10 nominally independent variants from the discovery trans - ancestral (Trans.) meta - analysis and the discovery meta - analyses in African (AA) and European (EU) ancestry cohorts, respectively. Independent variants are identified based on clumping for LD (pairwise r^2^ < 0.1) in 1000 Genomes Project Phase 3 data^25^. EU results are clumped using European (EUR) ancestry reference samples, AA results are clumped using African ancestry reference samples from the American Southwest (ASW), and trans - ancestral results are clumped using merged EUR and African ancestry (AFR) reference samples. Meta - analysis p - values and allele frequencies (Freq.) are reported from full discovery meta - analyses. Bold indicates genome - wide significant p - values (p < 5e - 8). Odds ratios (OR) and INFO scores are reported from the meta - analyses of the subset of unrelated individuals within each ancestry. Chromosome (CHR) and base pair (BP) position are reported for genome build hg19, with genes annotated by Ensembl VEP^62^. Allele frequency and OR are given with respect to allele 1 (A1).

Dashed red reference line indicates genome - wide significance (p < 5E - 8). Results are from the discovery meta - analysis of all cohorts (14,904 cases, 37,994 controls) under a fixed effects model weighted by effective sample size.

Results of meta - analysis with effective sample size weights for the *ADH1B* locus in (A) all cohorts, (B) AA cohorts, and (C) EU cohorts. Red reference line indicates the genome - wide significance threshold (P < 5e - 8). Within ancestry, colored points reflect the degree of LD (pairwise r^2^) to the index variant (indicated by a purple diamond) in 1000 Genomes Project reference data^25^ for individuals of (B) African or (C) European ancestry, respectively. No reference LD panel exists for the trans - ancestral sample (A).

**Figure 3:**
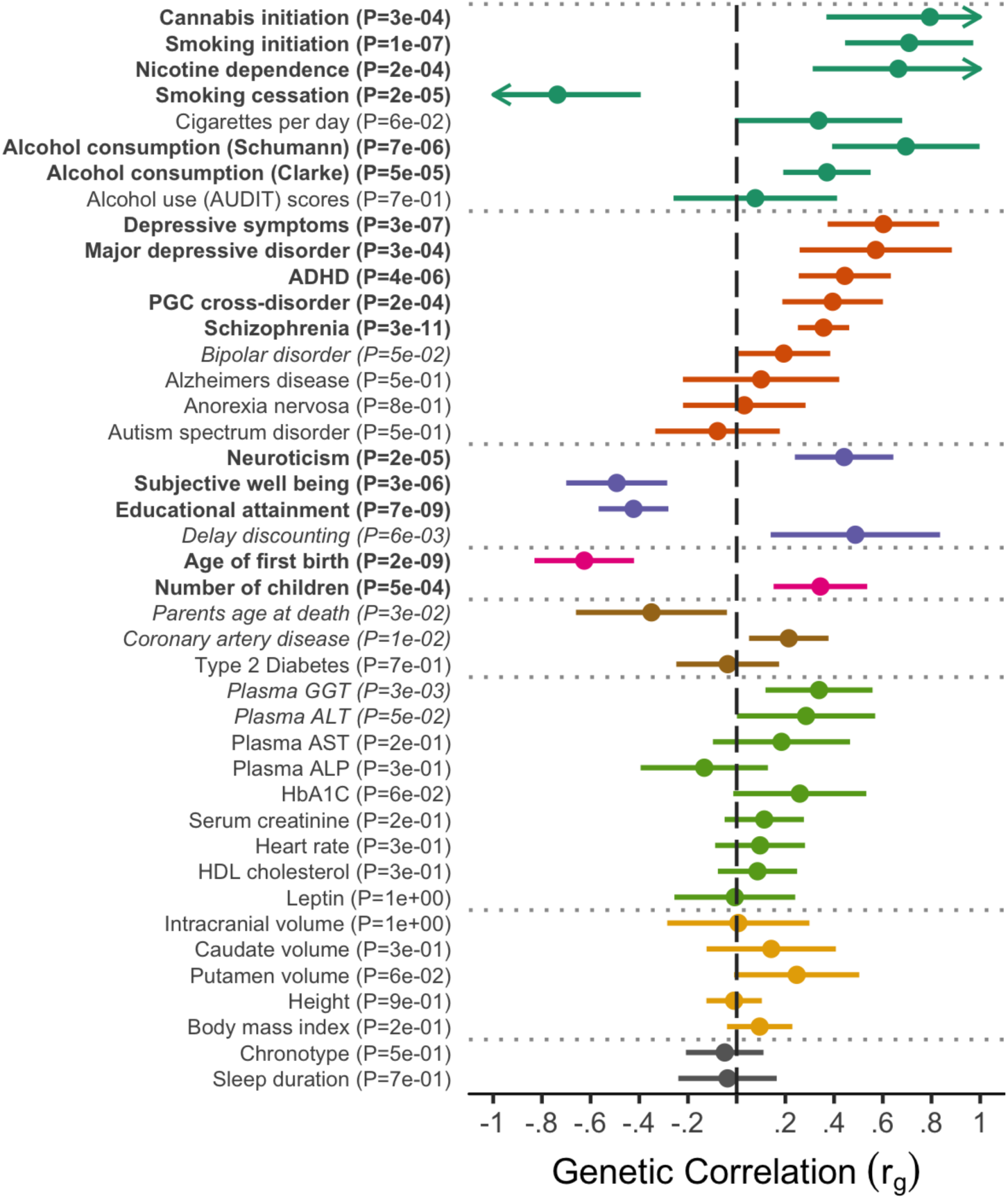
Genetic correlations between 42 traits and alcohol dependence in Europeans.

Genetic correlation results from LD score regression with the meta - analysis of AD in unrelated EU individuals (10,206 cases, 28,480 controls). After Bonferroni correction, significant correlations are observed with 16 traits and disorders (p < 1.2E - 3); bold), with nominally significant results for 6 additional traits and disorders (p < .05; italics). Error bars indicate 95% confidence intervals, with arrows indicating intervals extending above 1 or below - 1. Phenotypes are organized by research domain.

## Ethics statement

This study was approved by the institutional review board (IRB) of Washington University in St. Louis (Human Research Protection Office; number 201512068). Each contributing cohort obtained informed consent from their participants and received ethics approvals of their study protocols from their respective review boards in accordance with applicable regulations.

## Contributors

The 23andMe research team includes Michelle Agee, Babak Alipanahi, Adam Auton, Robert K. Bell, Katarzyna Bryc, Sarah L. Elson, Pierre Fontanillas, Nicholas A. Furlotte, David A. Hinds, Karen E. Huber, Aaron Kleinman, Nadia K. Litterman, Jennifer C. McCreight, Matthew H. McIntyre, Joanna L. Mountain, Elizabeth S. Noblin, Carrie A.M. Northover, Steven J. Pitts, J. Fah Sathirapongsasuti, Olga V. Sazonova, Janie F. Shelton, Suyash Shringarpure, Chao Tian, Joyce Y. Tung, Vladimir Vacic, and Catherine H. Wilson

## Data availability

Summary statistics from the genome - wide meta - analyses will be made available on the Psychiatric Genomics Consortium’s downloads page (http://www.med.unc.edu/pgc/results - and - downloads). Individual - level data from the genotyped cohorts and cohort - level summary statistics will be made available to researchers following an approved analysis proposal through the PGC Substance Use Disorder group with agreement of the cohort PIs; contact the corresponding authors for details. Cohort data is also available from dbGaP except where prohibited by IRB or European Union data restrictions (accession numbers to be available before publication).

## Code availability

Code for GWAS of case/control cohorts with ricopili is available at https://github.com/Nealelab/ricopili. Code for GWAS of family - based cohorts with picopili is available at https://github.com/Nealelab/picopili. Code for LD score regression analyses are available at https://github.com/bulik/ldsc. Effective sample size calculations were implemented using PLINK (http://www.cog-genomics.org/plink2), and GMMAT (https://content.sph.harvard.edu/xlin/software.html#gmmat) and geepack (https://cran.r-project.org/web/packages/geepack/index.html) in R (https://cran.r-project.org/); example code is available from the first author by request.

